# Light-Inducible Recombinases for Bacterial Optogenetics

**DOI:** 10.1101/786533

**Authors:** Michael B. Sheets, Wilson W. Wong, Mary J. Dunlop

**Affiliations:** Department of Biomedical Engineering, Boston University, Boston, MA 02215; Biological Design Center, Boston University, Boston, MA 02215

## Abstract

Optogenetic tools can provide direct and programmable control of gene expression. Light-inducible recombinases, in particular, offer a powerful method for achieving precise spatiotemporal control of DNA modification. However, to-date this technology has been largely limited to eukaryotic systems. Here, we develop optogenetic recombinases for *Escherichia coli* which activate in response to blue light. Our approach uses a split recombinase coupled with photodimers, where blue light brings the split protein together to form a functional recombinase. We tested both Cre and Flp recombinases, Vivid and Magnet photodimers, and alternative protein split sites in our analysis. The optimal configuration, Opto-Cre-Vvd, exhibits strong blue light-responsive excision and low ambient light sensitivity. For this system we characterize the effect of light intensity and the temporal dynamics of light-induced recombination. These tools expand the microbial optogenetic toolbox, offering the potential for precise control of DNA excision with light-inducible recombinases in bacteria.

## Introduction

Optogenetic tools enable novel applications for synthetic biology.^1–5^These tools typically use light to control expression of genes, often relying on light-dependent changes in protein state to control protein-protein interactions,^6,7^ promoter systems,^8,9^ and ion channels.^10^ Optogenetic systems offer many advantages over traditional chemical approaches for controlling gene expression due to the direct and programmable nature of light as an input. Using light instead of small molecules can give precise spatiotemporal control over regulation, and can circumvent the need to change media or otherwise disrupt the system to add or remove a chemical inducer. As light is easily programmable using electronics, optogenetic tools can also interface with dynamic computer-based control and feedback.^11,12^

Microbial optogenetic approaches have revealed a myriad of new applications that take advantage of the precise, programmable nature of light. As examples, light has been used to control expression of enzymes involved in biofuel synthesis^13,14^ and to regulate bacterial growth via metabolic control.^15,16^ In addition, it has been used to enable light-activated drug release from hydrogels^17^ and patterning of *Escherichia coli* onto multiple materials,^18^ indicative of the wide ranging potential of optogenetic approaches. At present, the current bacterial optogenetic toolset primarily includes two-component systems^8,19,20^ and split proteins.^7,21^

Recombinases are proteins that recognize specific 30-50 base pair (bp) sequences of DNA, and excise the “target” DNA between the sites along with one of the recognition sites. Their ability to manipulate DNA makes them particularly useful for complex cellular logic circuits and engineering gene circuits with memory.^22,23^ Light-inducible recombinases have been notably useful in mammalian systems^6,24,25^ and yeast.^26^ Having recombinases that are inducible at the protein-level can allow specific cells within a population to be targeted for recombination in response to spatial patterning of light, and there is no need to change media or wait for a chemical inducer to diffuse. Recombinase enzymes can be made light sensitive by splitting the gene into N-terminal and C-terminal fragments, and linking a sequence for a light-sensitive photodimer to each fragment. Upon light induction, the photodimer undergoes a conformational change that allows it to dimerize, bringing the two fragments together. This split-protein approach has been shown to work for both chemogenetic and optogenetic split-recombinases in eukaryotic systems.^6,25^ However, light-inducible recombinases have remained largely in the eukaryotic domain.

Here, we develop and optimize an optogenetic recombinase for *E. coli*. We focus primarily on split Cre linked to Vivid (Vvd) photodimers. Cre is a commonly used tyrosine recombinase from the P1 bacteriophage that excises DNA flanked by *loxP* sites.^27^ Vvd is derived from the fungus *Neurospora crassa*, and homodimerizes under blue light and separates in the dark.^28,29^ In developing our light-inducible recombinase we also explored Flp recombinase and Magnet photodimers,^30^ as well as multiple protein split sites within each recombinase. Here, we introduce an optimized design, which we denote Opto-Cre-Vvd, which excises target DNA completely in 2 hours. We also characterize sensitivity to ambient light exposure, the impact of light intensity, and the response time of the system.

## Results

To make Cre light sensitive, we split it into N-terminal (nCre) and C-terminal (cCre) fragments, with Vvd photodimers attached to the internal end of each fragment (Fig. 1a). When exposed to blue light, Vvd changes conformation to allow dimerization, bringing the Cre fragments together.

**Figure 1:**
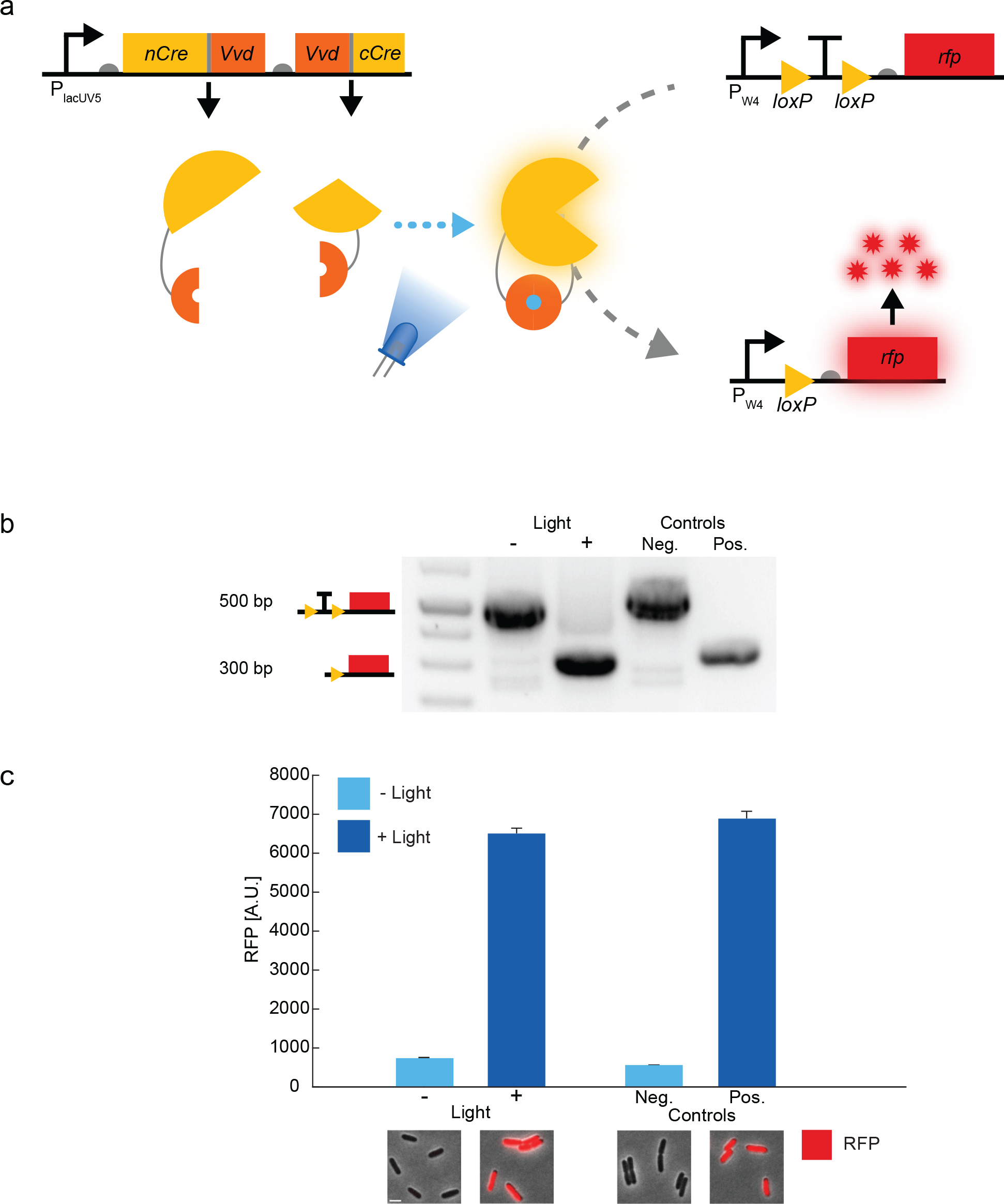
Light-inducible recombination in *E. coli*. **(a)** Split Cre fragments are linked to Vvd photodimers and expressed under the control of an IPTG-inducible promoter (P_lacUV5_). When exposed to blue light, Vvd dimerizes, forming functional Cre protein. Cre can then act on the reporter plasmid, excising the *loxP*-flanked transcription terminator and allowing expression of RFP. RFP is under the control of a constitutive promoter (P_W4_). **(b)** Gel electrophoresis images showing DNA excision. PCR of the reporter region containing *loxP*-flanked terminator shows a 500 bp band when full terminator is intact, and 300 bp band after recombination. Negative control contains cells with the reporter plasmid alone (terminator upstream of *rfp*); positive control contains cells with recombinase and a pre-cut reporter plasmid (no terminator upstream of *rfp*). **(c)** Single-cell fluorescence microscopy showing RFP expression for cells with and without light exposure. Insets below show representative cell images (scale bar = 2 μm). Error bars show standard error around the mean (n ≈ 300 cells per sample).

Cre excises DNA fragments between *loxP* sites that are oriented in the same direction. We used this to develop a reporter for the efficiency of our recombinase constructs. We placed a transcription terminator flanked by *loxP* sites between the gene for red fluorescent protein (*rfp*) and a constitutive promoter. In the absence of Cre, the terminator prevents transcription of *rfp*. Functional Cre excises the terminator, leading to RFP production (Fig. 1a). To perform these tests, we used a light plate apparatus (LPA).^31^ We exposed samples to 465 nm blue light using LEDs for one hour and then took samples for polymerase chain reaction (PCR) immediately following light exposure. To verify that the recombinase was excising the target DNA properly, we first used PCR to check the length of the plasmid region containing the *loxP*-flanked terminator with and without exposure to light. We used a forward primer upstream of the promoter, and a reverse primer in the *rfp* gene to amplify the region, which is approximately 500 bp if the terminator and both *loxP* sites are intact, and 300 bp when the terminator is removed by recombination (Fig. 1b). As a negative control, we used cells with the reporter but no recombinase. As a positive control, we used a strain containing recombinase and the reporter plasmid with the terminator excised. In addition, using microscopy we confirmed that after exposure to blue light, cultures showed a clear increase in RFP (Fig. 1c). For the microscopy experiments we refreshed cultures overnight to allow full RFP expression and maturation after recombination.

When developing the photoactivatable split recombinase, we considered several variants on the design, including different recombinase enzymes, alternative photodimers, and multiple protein split site locations. First, we tested two widely-used recombinases, Cre and Flp (Fig. 2a). Flp is a tyrosine recombinase originally native to *Saccharomyces cerevisiae*,^32^ and like Cre has been used as a split photo-activatable recombinase in mammalian systems.^24,25^ We tested each recombinase using previously established split sites with two photodimer options, Vvd and Magnets. In contrast to the blue light-sensitive homodimer Vvd, Magnets are engineered heterodimer Vvd variants with separate positively-charged and negatively-charged dimer interfaces.^30^ We worked with these photodimers due to their prevalence in split protein engineering.^6,7,21,24,25,33^ We found that Cre, especially when paired with Vvd, showed substantially improved activation relative to Flp when exposed to blue light (Fig. 2a).

**Figure 2:**
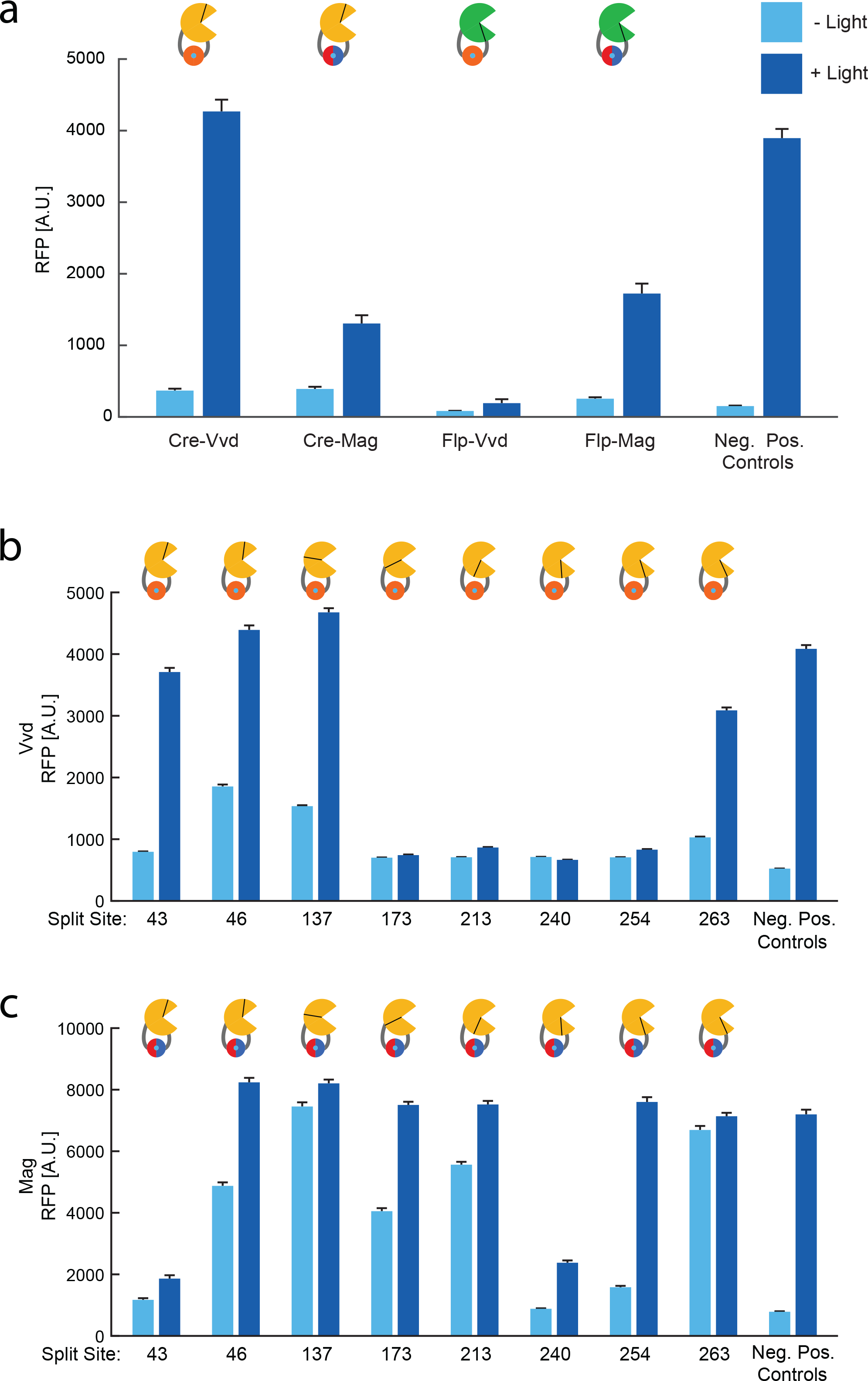
Comparison of optogenetic recombinase protein variants. **(a)** RFP reporter output with and without light exposure for Cre and Flp recombinases, each with Vvd and Magnet photodimers. Split sites used are Cre with nCre length of 43 AA, Flp with nFlp length of 374 AA. **(b)** Assay of split sites tested for Cre-Vvd and **(c)** Cre-Mag. Numbers shown for split sites on the x-axis are the length of nCre. All figure data obtained using fluorescence microscopy. Error bars show standard error around the mean (n ≈ 750 cells per sample).

Focusing on Cre recombinase with the Vvd photodimer, we next tested several variants where the protein was split at different locations (Fig. 2b). We considered sites reported in the literature, structurally-predicted sites, and algorithmically-derived sites (Table 1). Literature derived sites included Cre 43,^6^ and Cre 213, 240, 254.^25^ We also used the SPELL algorithm to determine novel potential split sites.^34^ We ran SPELL on the Cre structure PDB 3MGV,^35^ which led to predictions for Cre 46 and Cre 137. We also found structurally-informed sites for Cre by analyzing the PDB structure 3MGV in PyMol.^36^ Using this crystal-derived structure, we assessed B-factor to select flexible regions within the protein,^37^ and chose split sites between flexible amino acids such as glycine and serine.^38^ This method led us to Cre 172 and 263.

**Table 1.**
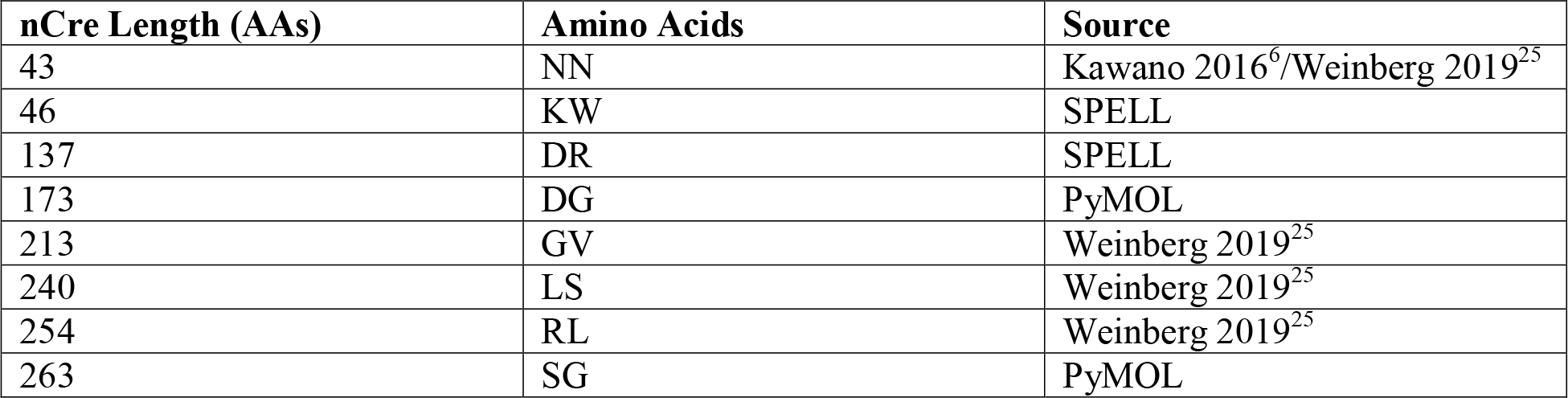
Cre recombinase split sites used in this study, given as lengths of the nCre fragment (from methionine at position 1 to the split site), the amino acids on either side of the split, and the source of the split. Note that we use truncated Cre from Kawano (2016), which starts at AA 18 of native Cre.

| nCre Length (AAs) | Amino Acids | Source |
| --- | --- | --- |
| 43 | NN | Kawano 2016<sup>6</sup>/Weinberg 2019<sup>25</sup> |
| 46 | KW | SPELL |
| 137 | DR | SPELL |
| 173 | DG | PyMOL |
| 213 | GV | Weinberg 2019<sup>25</sup> |
| 240 | LS | Weinberg 2019<sup>25</sup> |
| 254 | RL | Weinberg 2019<sup>25</sup> |
| 263 | SG | PyMOL |

We observed split site-dependent variation in both the level of activation with light and in recombination in the absence of light. Some sites showed almost no activation with light (173, 213, 240, 254). Others sites showed high activation with light exposure, but also increased activation in the absence of light (46, 137). From this screen, we found Cre-Vvd split after AA 43 to be our best candidate, as it showed a good fold change in RFP expression in response to light and minimal activation without light. We also tested each split site using the Magnet photodimers (Fig. 2c), and observed varied activation at different split sites. Although there were commonalities, not all split sites behaved consistently with both types of photodimers. Overall, we found that Magnets were more likely to strongly activate, but also had much higher activation without light than Vvd. This may be due in part to Vvd’s ability to form homodimers, as “incorrect” dimer pairs containing two nCre or two cCre fragments could help to lower formation of Cre in the absence of light. It is also notable that the dark-state expression seen here is higher than in the original mammalian PA-Cre.^6^ Tests of five literature-derived split sites for Flp-Mag showed similar split site-dependent results, but were ultimately inferior to the Cre variants (Fig. S1). Due to its high fold change we chose to use Cre-Vvd 43, which we denote Opto-Cre-Vvd, for further characterization.

An important practical experimental consideration for light inducible recombinases is their sensitivity to ambient light. Therefore, we next tested how Opto-Cre-Vvd performed with 5 minutes of ambient light exposure. We chose this duration to mirror conditions that might be experienced in a setting where plates are temporarily removed from darkness, such as would be necessary to transfer cultures from growth conditions to flow cytometry or microscopy assays. We found that Opto-Cre-Vvd showed minimal sensitivity to short duration exposure to ambient light (Fig. 3a).

**Figure 3:**
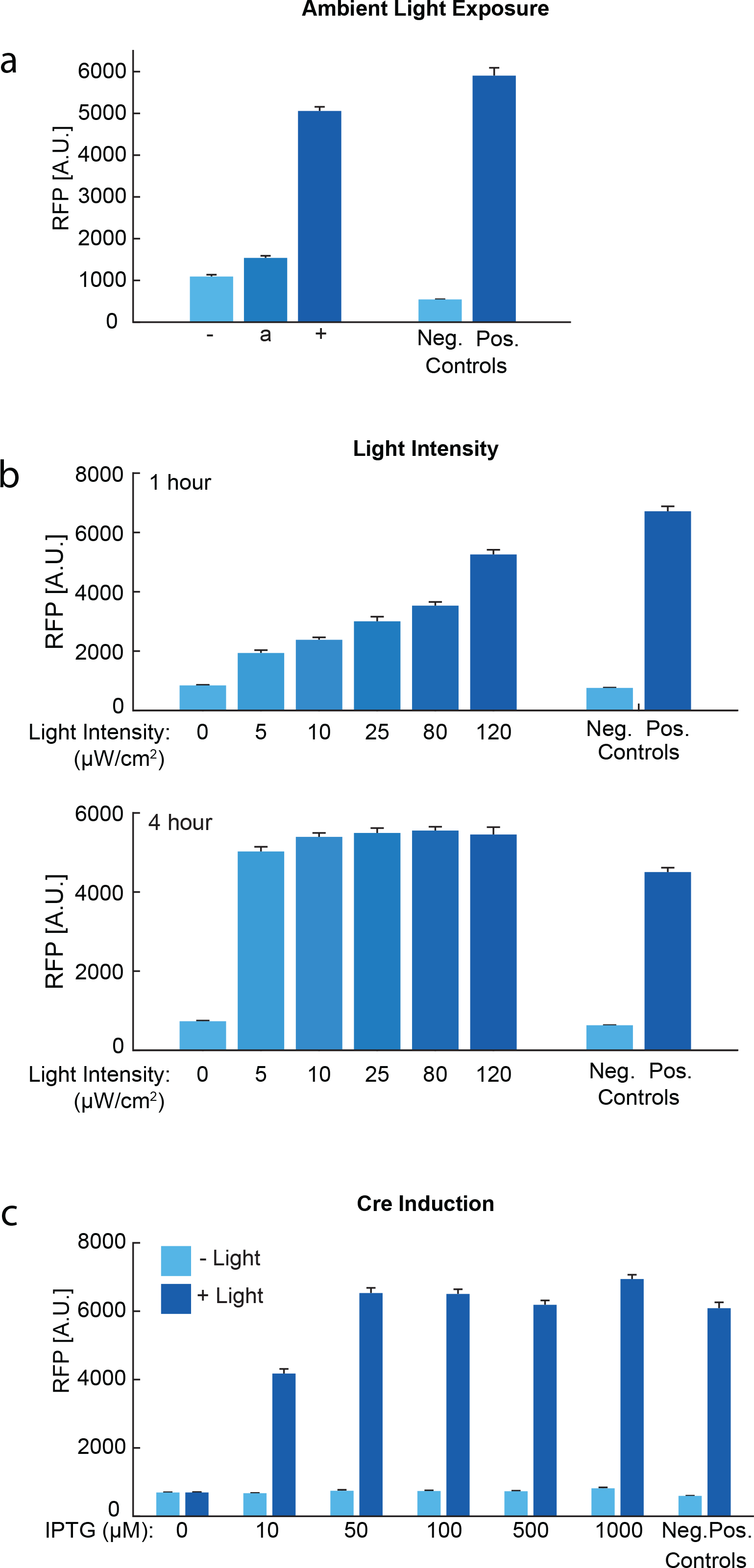
Characterization of Opto-Cre-Vvd. **(a)** RFP reporter output for Opto-Cre-Vvd without light (−), with a 5 minute exposure to ambient light (a), and with full exposure to blue light (+). **(b)** Effect of blue light intensity on Opto-Cre-Vvd activation when exposed for 1 or 4 hours. **(c)** Effect of IPTG induction levels for Opto-Cre-Vvd. In all cases, error bars show standard error around the mean from microscopy data (n ≈ 350 cells per sample).

Next, we tested experimental parameters for Opto-Cre-Vvd, including the light intensity used for induction, the timing of light exposure, and concentration of IPTG for recombinase induction. When optimizing split sites and photodimer variants, we used a blue light intensity that corresponded to the maximum value accessible with the LEDs used in the LPA (120 μW/cm^2^) to minimize excision times (Fig. 3b). Using microscopy, we observed no discernable in differences cell morphology with and without light exposure in these conditions, suggesting that phototoxicity effects were minimal with this exposure level (Fig. S2). However, we found that even at much lower light intensities, we observed complete excision when samples were exposed to blue light for a longer time (Fig. 3b). Cultures exposed to lower intensities of light showed partial excision after 1 hour, while cultures exposed to high intensity light showed near-complete excision. When we exposed cultures to constant blue light for 4 hours, we found that all intensities of light yielded comparable high levels of excision (Fig. 3b).

In our design, the split Cre fragments are under the control of a lacUV5 promoter to prevent excision of the target DNA prior to induction and subsequent light exposure. We found that inducing with IPTG concentrations above 50 μM for two hours prior to light exposure was sufficient to induce Cre for light activation (Fig. 3c). We used 100 μM IPTG as a standard value, which remains solidly above the induction threshold for our experiments.

We were also interested in exploring the duration of light exposure that cells need to induce full RFP expression (Fig. 4). To test this, we exposed separate cultures to light for 5 minutes, 30 minutes, 1 hour, 2 hours, 4 hours, or 8 hours. Cultures exposed to light for less than 8 hours were kept in the dark following light exposure for the remainder of the time. At the end of the 8 hour period, we refreshed all cultures and grew them for two hours without light to allow for protein maturation and then assessed transcription terminator excision both genotypically and phenotypically. Genotypic excision was assessed by PCR and gel electrophoresis (Fig. 4a). Phenotypic results were assessed by microscopy (Fig. 4b,c) and spotting on agar (Fig. 4d). We observed general agreement between all characterization methods. Opto-Cre-Vvd shows substantial RFP expression within 1 hour, and RFP values comparable to the positive control, indicative of near-complete activation, by 2 hours.

**Figure 4:**
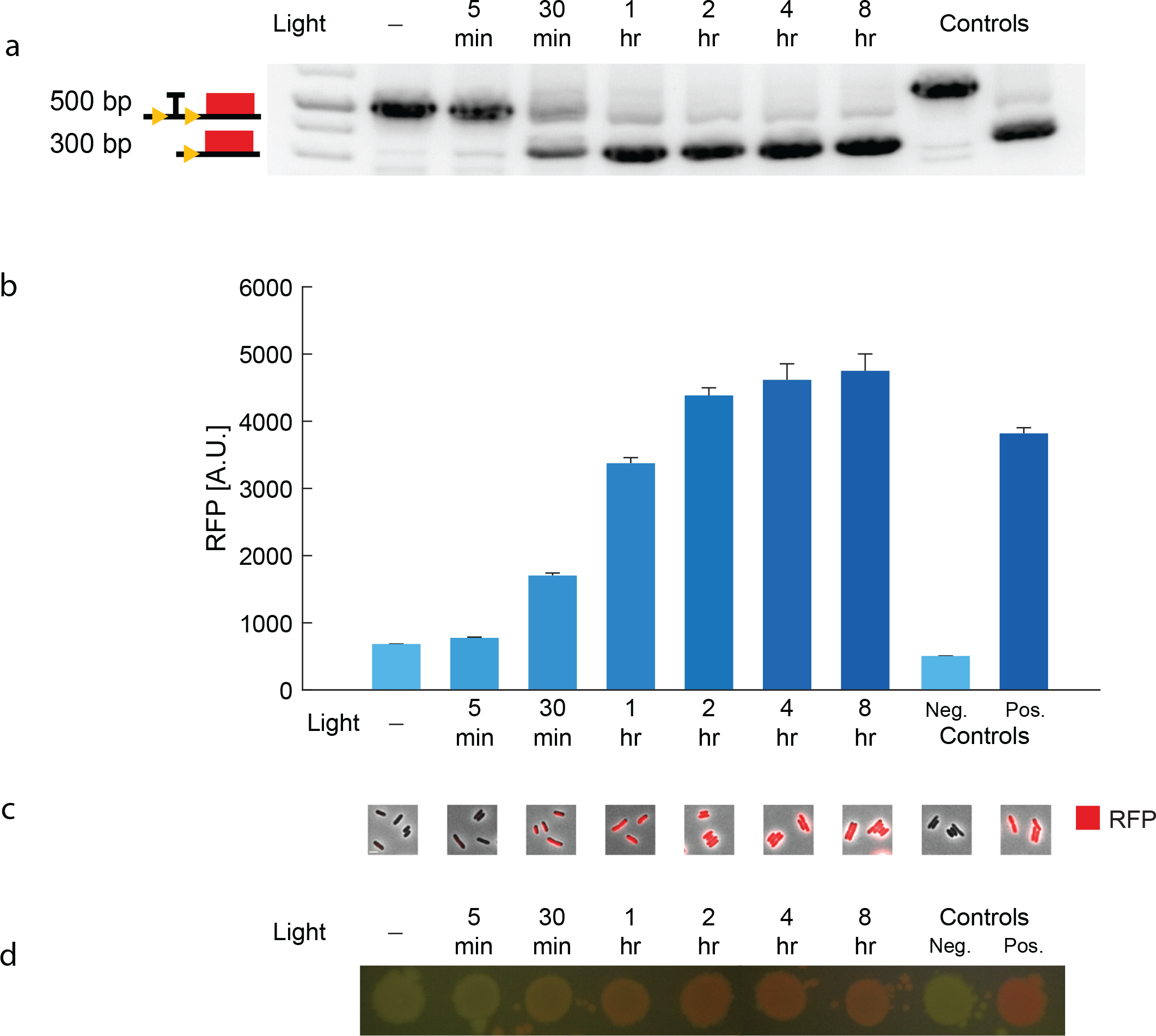
Light exposure duration necessary to induce excision by Opto-Cre-Vvd. **(a)** DNA gel image showing reporter bands with and without transcription terminator excision, **(b)** single-cell fluorescence microscopy averages of RFP values, **(c)** representative microscopy images (scale bar = 2 μm), and **(d)** samples of culture spotted on agar plates of Opto-Cre-Vvd exposed to different durations of blue light. Error bars show standard error around the mean (n ≈ 750 cells per sample).

## Discussion

We have developed, optimized, and characterized a light-inducible recombinase for *E. coli*. We found that both Cre and Flp associated with either Vvd or Magnet photodimers have the potential for photo-activatable recombination. However, split site location and the recombinase-photodimer pairing impact efficacy. Our most promising candidate, Opto-Cre-Vvd, exhibits blue light-dependent excision and low sensitivity to ambient light. We also found that Opto-Cre-Vvd shows activation at both low and high light intensities, but at different timescales. The construct can cut completely within 2 hours, which is comparable to the timeframe observed for mammalian photoactivatable Cre^6^ and existing Magnet-based split proteins in *E. coli*.^7^

A well-characterized recombinase tailored to *E. coli* is a powerful new tool for bacterial optogenetics. Moving forward, this offers expanded potential for interfacing engineered cells with computational control via light. Applications include the ability to target sub-populations of cells, real-time genetic modifications, or experiments where small molecule inducers are impractical due to crosstalk. A future version of this system may also be useful for changing genetic state during biomolecule production, such as in metabolic engineering applications, as light is an inexpensive inducer. Cre could also be used to activate or inactivate multiple genes. Along these lines, future extensions to this system may involve development of orthogonal light-inducible recombinases for bacteria. Alternate photodimer systems that are sensitive to other wavelengths of light could also be used to multiplex the approach.^39^ As an immediate application, Opto-Cre-Vvd can be used as-is to perform gene knock outs or insertions in real time and is compatible with plate-based or microscopy platforms. These light-inducible recombinases expand the optogenetic methods available for bacteria and have great potential for the design of novel synthetic circuits.

## Methods

### Strains and Plasmids

Expression studies use *E. coli* strain MG1655. All recombinase constructs use a plasmid with a high-copy ColE1 origin and an ampicillin resistance cassette where the recombinase genes are under the control of an IPTG-inducible lacUV5 promoter, derived from the pBbE5a BioBrick plasmid.^40^ All reporter constructs use a medium-copy p15A origin plasmid with a kanamycin resistance cassette and the gene for red fluorescent protein (mRFP1)^41^ under the control of a constitutive, medium-strength promoter (denoted P_W4_), which is modified from the phage T7 A1 promoter: TTATCAAAAAGAGTATTG<u>CA</u>TTAAAGTCTAACCTATAGGA<u>AT</u>CTTACAGCCATCGAGAGGGACACGGCGAA (underline indicates mutations from original T7 A1 promoter).^42^ Plasmids were constructed using the Gibson assembly method.^43^ Primer data can be found in Table S1.

Original Cre and Magnet heterodimer plasmids are from Weinberg (2019).^25^ Original Flp gene sequence was derived from the pCP20 plasmid from Datsenko & Wanner (2000).^44^ We obtained the Vivid homodimers from AddGene plasmid #58689 (mV-NcVV-LOV_231) deposited by Harald Janovjak.^29^ We express Cre split with a photodimer pair as an operon (Fig 1a). The N-terminal fragment of Cre (nCre) is followed by a 10 AA glycine-serine linker and a photodimer. A separate RBS is used to express the second photodimer linked by a 10 AA glycine-serine linker to the C-terminal fragment of Cre (cCre).

Plasmids from this study and their sequences are available on AddGene (https://www.addgene.org/Mary_Dunlop/).

### Cre Split Site Selection

Split sites for Cre were selected using three methods: chosen from the literature, using the first two optimal choices from the SPELL algorithm (https://dokhlab.med.psu.edu/spell/)^34^ based on the PDB structure 3MGV^35^, or by using PyMOL (https://pymol.org/2/) on 3MGV and selecting two sites around glycine and serine AAs in regions with high B-factor values. Further information and source of each split site for Cre can be found in Table 1. Primers used to make each split are listed in Table S1. Split Cre variants were made by amplifying from plasmids containing Cre without a photodimer from the location of the split site with overhangs for the linker sites. In parallel, we amplified the photodimer and linker inserts and combined via Gibson assembly.^43^ Split sites for Flp were chosen from the literature and cloned in a similar fashion; their information can be found in Table S2.

### Light Exposure Assays

Strains were grown overnight from a single colony in LB medium containing 100 μg/mL carbenicillin and 30 μg/mL kanamycin for plasmid maintenance. The next day, cultures were refreshed 1:100 in selective LB and induced for 2 hours with 100 μM IPTG unless otherwise noted. Blue light exposure was performed using a LPA,^31^ with two 465 nM wavelength LEDs per well (ThorLabs LED465E), outputting a total of 120 μW/cm^2^ Unless otherwise noted, cultures were exposed to blue light for 1 hour. After exposure, samples were prepared for analysis by PCR to check for target excision by gel electrophoresis, and refreshed in selective LB medium without IPTG overnight. In light intensity experiments, cultures were exposed to light for 4 hours total, with intermediate samples taken at 1 hour for characterization.

All liquid cultures through the experiment were grown at 37°C with 220 rpm shaking. Note that agar plates with the Flp recombinase and reporter were also kept at 37°C at all times, as we observed substantial activation even without light when stored at 4°C. After transformation, all cultures were kept in the dark throughout the entire experiment with the exception of blue or ambient light exposure periods. For ambient light exposure, samples were exposed to lab lighting in a shaking incubator for 5 minutes after induction, and then kept in the dark for the remainder of the experiment.

### Recombinase Efficiency Characterization

Target DNA excision via the recombinase was measured genetically by amplifying the promoter region of the reporter using PCR. We used a forward primer ~200bp upstream of the first *loxP* site (ATCTTCCCCATCGGTGATGTCG) and a reverse primer ~100bp downstream of the second *loxP* site (GACGACCTTCACCTTCACCTT) to check for differences in band length before and after recombination.

In addition to the PCR-based measurements, efficiency was also measured visually by imaging plated samples with a mobile phone camera using standard settings (Samsung Galaxy Note 8) on a Blue LED transilluminator through the attached orange filter (IO Rodeo).

### Microscopy and Image Analysis

Post light-exposure samples were refreshed overnight in LB medium with 100 μg/mL carbenicillin and 30 μg/mL kanamycin for plasmid maintenance to allow full RFP expression and maturation. Before imaging, samples were refreshed for 2 hours in 1:100 in MGC medium (M9 salts supplemented with 2 mM MgSO_4_, 0.2% glycerol, 0.01% casamino acids, 0.15 μg/ml biotin, and 1.5 μM thiamine). Samples were then placed on 1.5% low melting agarose pads made with MGC medium. Cells were imaged at 100x using a Nikon Ti-E microscope. Images were segmented and analyzed using the SuperSegger software^45^ and custom Matlab analysis scripts.

## Acknowledgements

We thank Elliot Tague and John Ngo for their input on split site selection; Ben Weinberg, Armin Baumschlager, and Mustafa Khammash for helpful discussions on experimental design; and Nathan Tague for early work on light-inducible Flp. Nadia Sampaio and Nathan Tague provided helpful comments on the manuscript. This work was supported by NIH grants R21AI137843 (MJD and WWW) and R01AI102922 (MJD). WWW acknowledges funding from the NIH Director’s New Innovator Award (1DP2CA186574), NSF Expeditions in Computing (1522074), NSF CAREER (162457), NSF BBSRC (1614642), NSF EAGER (1645169), and Boston University College of Engineering Dean’s Catalyst Award. MBS received support through the NIH training grant T32 EB006359.

## Author Contributions

MBS and MJD conceived and designed the experiments. MBS performed the experiments and analyzed the data. WWW provided insight on experimental design and optimization. MBS and MJD wrote the manuscript with input from WWW.

## Abbreviations

Vvd: Vivid
bp: base pair
nCre: N-terminal fragment of Cre recombinase
cCre: C-terminal fragment of Cre recombinase
RFP: Red Fluorescent Protein
PCR: Polymerase Chain Reaction
AA: Amino Acid
LPA: Light Plate Apparatus
SPELL: Split Protein Reassembly by Ligand or Light

