## Supporting Information for "Light-Inducible Recombinases for Bacterial Optogenetics"

**Table of Contents**

Supplementary Figures

Supplementary Tables

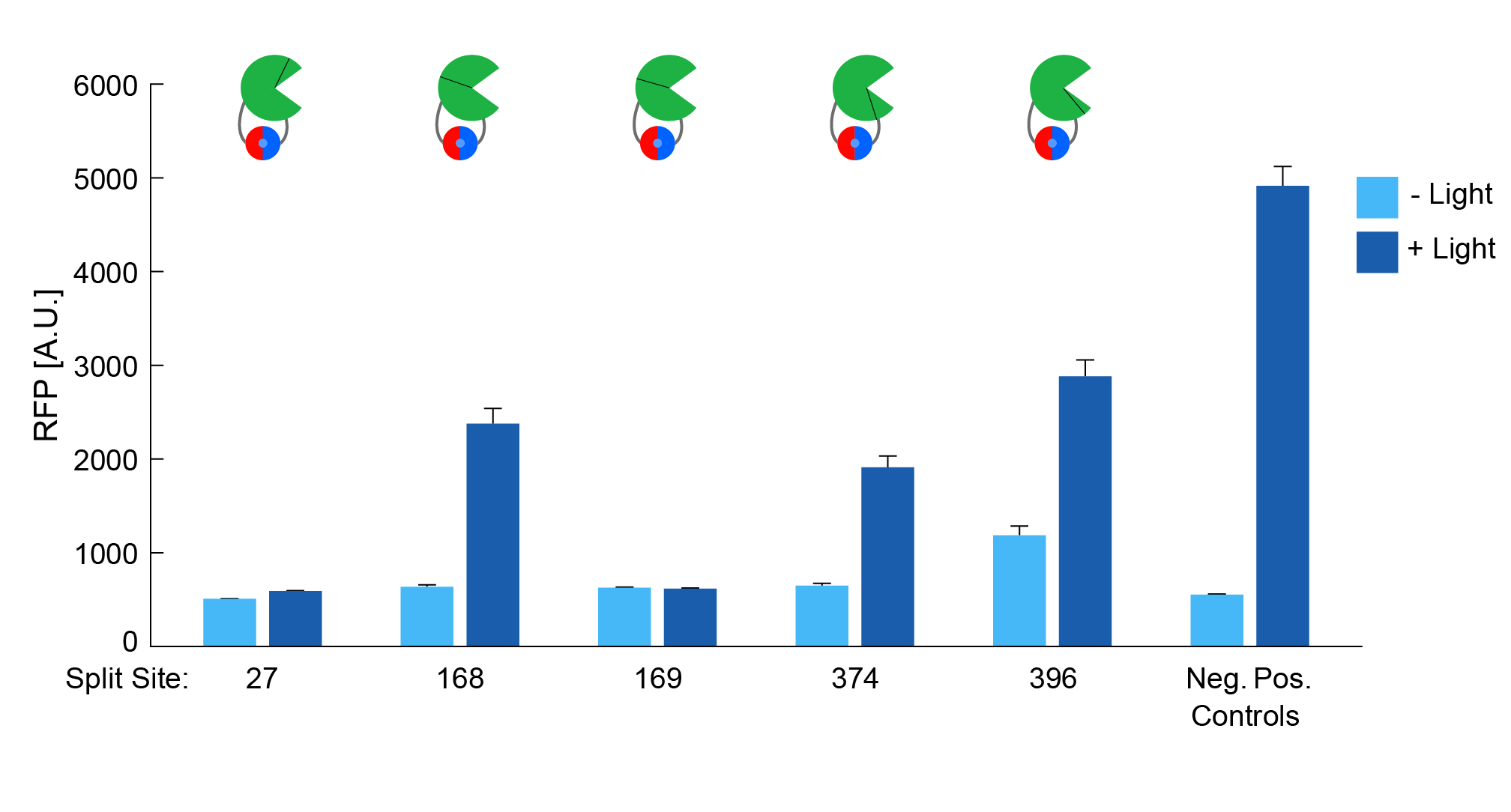

**Figure S1**: Split sites for Flp-Mag. Numbers shown for split sites on the x-axis are the length of nFlp. Error bars show standard error around the mean for microscopy data (average n ≈ 100 cells per sample).

**­­**

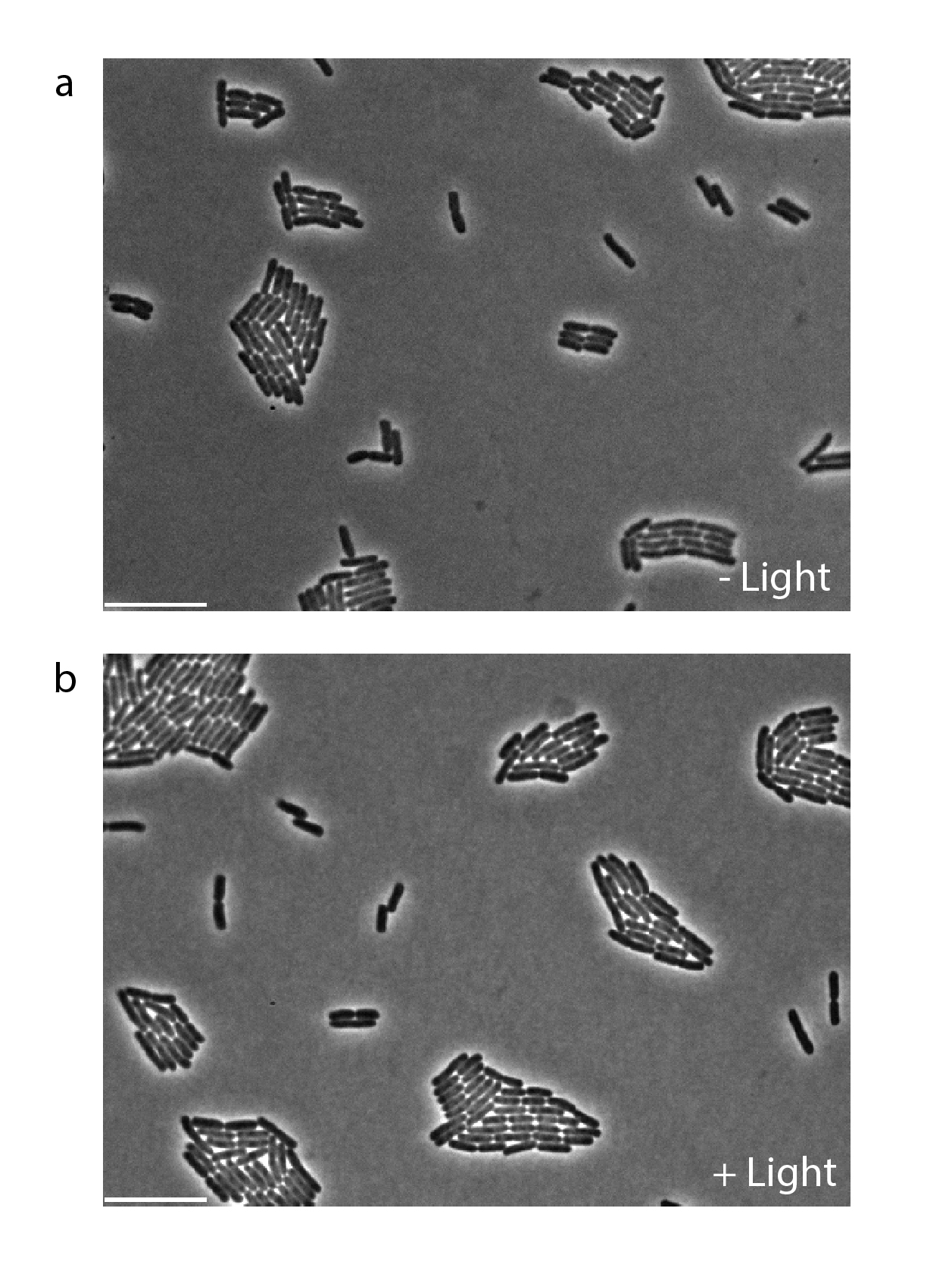

**Figure S2**: Single-cell phase contrast microscopy of fully induced Opto-Cre-Vvd cells **(a)** without blue light exposure, and **(b)** after 1h of 120 μW/cm^2^ blue light exposure. Scale bar = 10 μm.

**Table S1.** Primers used for plasmid assembly of split site constructs. Overhang regions are shown in lowercase.

| Mag/Vvd + Linkers Insert F | GGGTCTGGCTCCGGATC |
| --- | --- |
| Mag/Vvd + Linkers Insert R | CGGAGGTTCAGGAGGTTCA |
| Cre-43-F | ccggaggttcaggaggttcaAACAGGAAATGGTTCCCTGC |
| Cre-43-R | cttgatccggagccagacccGCCTGGTGCAAGCTGAAC |
| Cre-46-F | ccggaggttcaggaggttcaTGGTTCCCTGCTGAACCT |
| Cre-46-R | cttgatccggagccagacccTTTCCTGTTGTTCAGCTTGCA |
| Cre-137-F | ccggaggttcaggaggttcaAGATGCCAGGACATCAGGA |
| Cre-137-R | cttgatccggagccagacccGTCAGAGTTCTCCATCAGGGA |
| Cre-173-F | ccggaggttcaggaggttcaGGTGGGAGAATGCTGATCC |
| Cre-173-R | cttgatccggagccagacccATCGGTGCGGGAGATGT |
| Cre-213-F | ccggaggttcaggaggttcaGTGGCTGATGACCCCAAC |
| Cre-213-R | cttgatccggagccagacccACCAGACACAGAGATCCATCT |
| Cre-240-F | ccggaggttcaggaggttcaTCCACCCGGGCCCT |
| Cre-240-R | cttgatccggagccagacccCAGTTGGGAGGTGGCAGA |
| Cre-254-F | ccggaggttcaggaggttcaCTGATCTATGGTGCCAAGGATG |
| Cre-254-R | cttgatccggagccagacccGCGGTGGGTGGCCTC |
| Cre-263-F | ccggaggttcaggaggttcaGGGCAGAGATACCTGGCC |
| Cre-263-R | cttgatccggagccagacccTGGTGCCAAGGATGACTCT |

**Table S2**. Flp recombinase split sites used, listed as lengths of the nFlp fragment (from methionine at position 1 to split site), the amino acids on either side of the split, and the source of the split.

| **nFlp Length (AAs)** | **Amino Acids** | **Source** |
| --- | --- | --- |
| 27 | SG | Jung 2019^1^/Weinberg 2019^2^ |
| 168 | TS | Weinberg 2019^2^ |
| 169 | SR | Jung 2019^1^ |
| 374 | LK | Weinberg 2019^2^ |
| 396 | GS | Weinberg 2019^2^ |
